# Shear force enhances adhesion of *Pseudomonas aeruginosa* by counteracting pilus-driven surface departure

**DOI:** 10.1101/2023.05.08.539440

**Authors:** Jessica-Jae S. Palalay, Ahmet N. Simsek, Benedikt Sabass, Joseph E. Sanfilippo

## Abstract

Fluid flow is thought to prevent bacterial adhesion, but some bacteria use adhesins with catch bond properties to enhance adhesion under high shear forces. However, many studies on bacterial adhesion either neglect the influence of shear force or use shear forces that are not typically found in natural systems. In this study, we use microfluidics and single-cell imaging to examine how the human pathogen *Pseudomonas aeruginosa* interacts with surfaces when exposed to shear forces typically found in the human body (0.1 pN to 10 pN). Through cell tracking, we demonstrate that the angle between the cell and the surface predicts if a cell will depart the surface. We discover that at lower shear forces, type IV pilus retraction tilts cells away from the surface, promoting surface departure. Conversely, we show that higher shear forces counterintuitively enhance adhesion by counteracting type IV pilus retraction-dependent cell tilting. Thus, our results reveal that *P. aeruginosa* exhibits behavior reminiscent of a catch bond, without having a specific adhesin that is enhanced by force. Instead, *P. aeruginosa* couples type IV pilus dynamics and cell geometry to tune adhesion to its mechanical environment, which likely provides a benefit in dynamic host environments.

## Introduction

Bacteria experience fluid flow in natural environments and during host colonization. As a result, many bacterial species have evolved elaborate mechanisms to colonize environments with high shear forces (1–6). For example, *Escherichia coli* uses the protein FimH to promote adhesion in the human urinary tract (6–8). FimH is remarkable as mediates strong adhesion and employs a catch bond, which becomes stronger when shear force is applied (8–10). *Staphylococcus aureus* also uses adhesins with catch bond properties (11–14), highlighting that force-enhanced adhesins are conserved throughout bacteria. *Pseudomonas aeruginosa* exhibits surface motility against the direction of flow (15), which provides an advantage in certain geometric contexts (3). *Caulobacter crescentus* colonizes flow-rich environments due in part to their curved-rod shape, which promotes the attachment of daughter cells to the surface in high flow conditions (16). Furthermore, bacterial quorum sensing is suppressed by flow (6, 17–19), demonstrating the complexity of cell-cell interactions in flowing environments. Together, these features reveal that bacteria are significantly impacted by flow and demonstrate the acute need to incorporate flow into bacterial experiments.

There are two important physical parameters associated with fluid flow: shear force and shear rate (20, 21). Shear force represents the force acting on cells that results in deformation or bending. Shear rate is a force-independent parameter related to the velocity of the fluid. While both shear force and shear rate depend on flow rate and microfluidic channel dimensions, only shear force depends on the viscosity of the solution (20). Thus, changing viscosity is an effective way to differentiate between the effects of shear force and shear rate (20). For example, an experiment with *E. coli* using varied viscosities led to the conclusion that FimH is a force-enhanced adhesin (21). In contrast, an experiment with *P. aeruginosa* using varied viscosities led to the conclusion that *froABCD* expression is triggered by shear rate (22). The conclusion that shear rate triggers *froABCD* expression guided subsequent work that demonstrated *froABCD* expression is regulated by the combined effects of flow and H_2_O_2_ transport (22). Together, research on FimH and *froABCD* highlights the power of changing viscosity to understand the biophysical and molecular mechanisms underlying flow-sensitive processes.

The study of bacterial adhesion has focused primarily on how bacteria arrive on a surface (2, 23–25). Bacterial surface arrival is dependent on multiple factors, including type IV pili (24–29). Type IV pili are dynamic filaments which extend and retract from the cell body and enable bacterial cell movement (26–33). The type IV pilus filament is made up of PilA monomers, while retraction of the filament is powered by motor proteins PilT and PilU (24, 30). In addition to their role in surface arrival, type IV pili have also been reported to promote surface departure of *P. aeruginosa* (32, 34) and *Vibrio cholerae* (33, 35). However, the examination of how type IV pili promote surface departure has not been well explored in fluid flow. One study (36) that did focus on flow revealed the counter-intuitive result that increasing flow enhanced the surface residence time of *P. aeruginosa*. However, the mechanism underlying flow-enhanced residence time remains unknown, highlighting the need for more research on bacterial surface departure in flow.

Here, we determine how the combined effects of shear force and type IV pili coordinate the surface departure of *P. aeruginosa*. Using microfluidics and single-cell imaging, we establish how type IV pili affect surface residence time and the surface orientation of cells in flow. We also utilize precise cell tracking to characterize how the presence of type IV pili change the adhesive properties of *P. aeruginosa* cells. By modulating solution viscosity, we quantitatively determine how shear force impacts surface departure in a host-relevant flow regime. Additionally, our data provide an explanation for the mechanism underlying flow-enhanced residence time. We propose that flow-enhanced residence time depends on shear force tilting cells toward the surface and restricting their motion. Collectively, our results highlight how biological and physical parameters combine to promote bacterial surface departure in host-relevant contexts.

## Results

To characterize how fluid flow affects *P. aeruginosa* surface interactions, we performed single-cell imaging of bacteria colonizing surfaces. To generate fluid flow, we custom-fabricated microfluidic devices, which were then connected to a precisely controlled syringe pump (Figure 1A). As surface arrival is the first step in bacterial surface colonization, we imaged and quantified wildtype *P. aeruginosa* PA14 cells arriving on a glass surface. For this experiment, microfluidic channels started with no cells and were imaged as cells were introduced into the channel in flow. Specifically, cells were loaded into a syringe at a mid-log concentration and were introduced at a shear rate (a measure of fluid flow) of 800 s^-1^. As cells flowed into the channel, many cells arrived on the surface and remained attached (Figure 1A). We quantified the cell arrival rate based on the number of cells that arrived and remained attached to the surface for at least 2 seconds (Figure 1B). Over the 60 second experiment, a significant number of wildtype cells arrived and began to accumulate on the surface.

**Figure 1:**
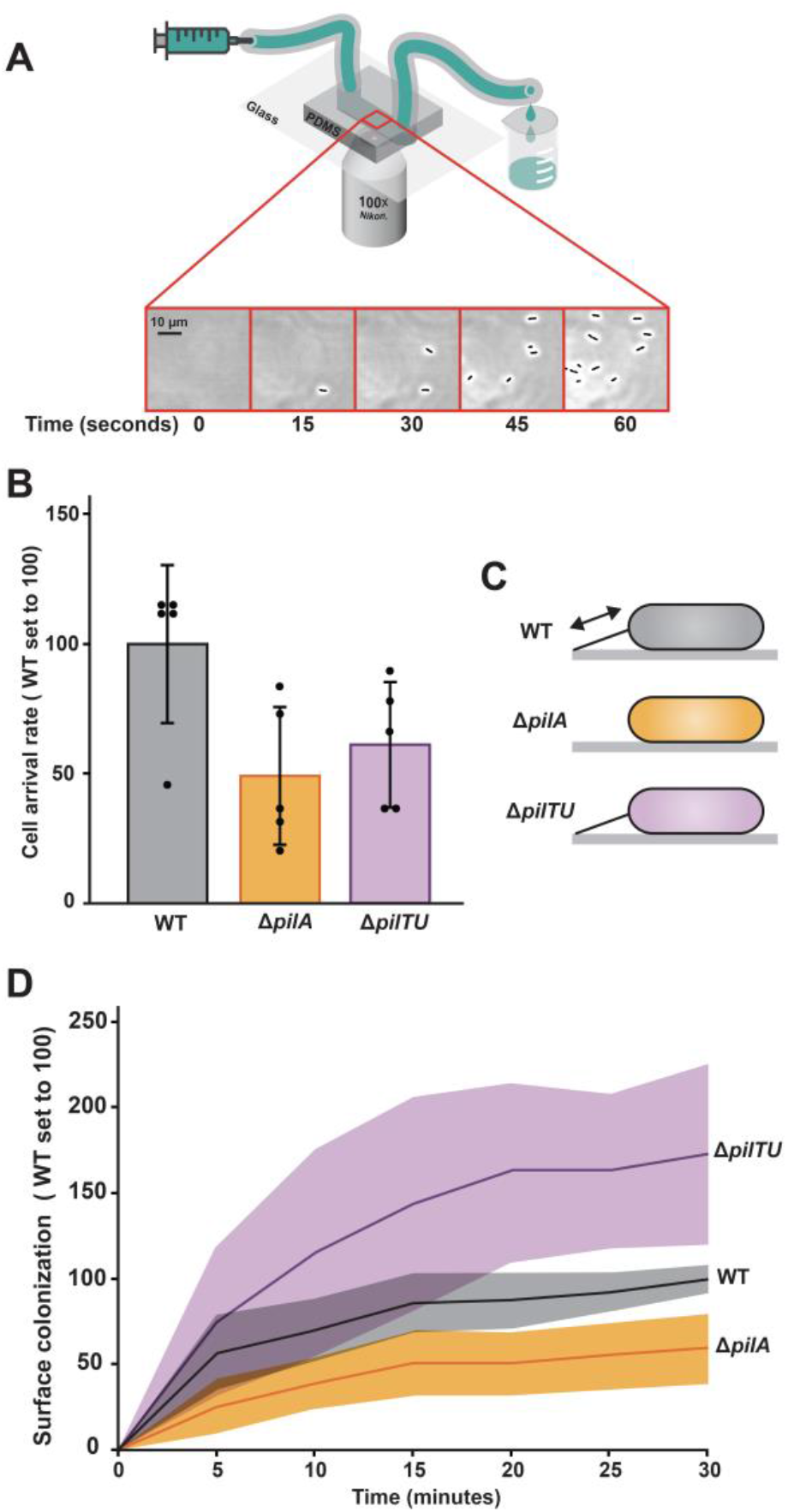
Type IV pili modulate *P. aeruginosa* surface colonization in flow. (A) Representation of microfluidic set-up used to observe cells throughout this study, reproduced with modification from (22) (Top). Microfluidic channels are made from polydimethylsiloxane (PDMS) and glass cover slips. Representative phase contrast images of bacterial cell colonization over 60 seconds (Bottom). Scale bar, 10 μm. (B) Cell arrival rate of wildtype (WT) (gray bar), Δ*pilA* (orange bar), and Δ*pilTU* (purple bar) cells at shear rate of 800 sec^-1^. Cell arrival rate was determined by how many cells landed on the surface and remained attached for at least 2 seconds. Quantification shows the average and standard deviation of five biological replicates. WT cell arrival rate was normalized to 100. WT and Δ*pilA* are statistically different with P = 0.03, while WT and Δ*pilTU* are statistically indistinguishable with P = 0.06. (C) Representation of WT, Δ*pilA*, and Δ*pilTU* cells. WT has dynamic pili (capable of extending and retracting), Δ*pilA* lacks pili, and Δ*pilTU* has pili present that typically lack dynamics. (D) Surface colonization of an empty channel with cells flowing into microfluidic device over a 30 minute time period. WT (black line and gray shading), Δ*pilA* (orange line and shading), and Δ*pilTU* (purple line and shading) cells were introduced at a shear rate of 800 sec^-1^. WT cell surface accumulation at 30 minutes is normalized to 100. Shading represents the standard deviation of three biological replicates.

How do type IV pili influence *P. aeruginosa* surface arrival? As type IV pili are known to affect attachment in many bacteria including *P. aeruginosa* (2, 23, 24, 37), we hypothesized that type IV pili would have an important role in surface arrival in our microfluidic assay. To test the role of type IV pili, we generated a Δ*pilA* mutant (which lacks pili) and a Δ*pilTU* mutant (which lacks typical pilus retraction) (Figure 1C). We measured surface arrivals in the Δ*pilA* and Δ*pilTU* strains and compared their surface arrival rate to wildtype (Figure 1B). The surface arrival rate of Δ*pilA* cells was approximately 50% lower than wildtype, supporting our hypothesis that type IV pili are important for surface arrival. In contrast, the surface arrival rate of Δ*pilTU* cells was not statistically different than wildtype. Together, these results suggest that the presence of type IV pili contribute to surface arrival.

To measure the effect of type IV pili on surface colonization, we flowed wildtype cells into empty microfluidic channels at a shear rate of 800 s^-1^ over 30 minutes (Figure 1D). Then, we imaged and quantified surface colonization, which we defined as the number of cells that accumulated on the surface. We observed that wildtype cells accumulated quickly for the first 15 minutes of the experiment, and then the total number of cells began to plateau (Figure 1D). To test the role of type IV pili on surface colonization, we repeated the experiment with Δ*pilA* and Δ*pilTU* strains. While Δ*pilA* cells colonized the surface approximately 50% less than wildtype, Δ*pilTU* cells colonized the surface approximately 75% more than wildtype. To confirm that the Δ*pilA* and Δ*pilTU* mutations were linked to the surface colonization phenotypes, we reintroduced *pilA* and *pilTU* into the genomes of the respective mutants and repeated our experiment. Reintroduction of *pilA* or *pilTU* returned the mutant phenotypes to wildtype levels, indicating that mutations in *pilA* and *pilTU* were causative of the surface colonization phenotypes (Figures S1, S2). While the reduction in colonization observed in the Δ*pilA* mutant can be explained by the reduction in surface arrival rate (Figure 1B), the mechanism underlying the enhancement of surface colonization in the Δ*pilTU* mutant is a mystery.

As surface colonization is affected by surface arrival and departure, the enhanced colonization phenotype of the Δ*pilTU* mutant could be explained by less surface departures. To test if Δ*pilTU* cells depart the surface less than wildtype, we forcibly removed cells from the surface using flow. We reasoned that if Δ*pilTU* cells experience less departures, they would be harder to remove from the surface with flow. Fluid flow exerts a shear stress on cells that is proportional to the shear rate and the solution viscosity (20) (Figure 2A). The shear force that cells experience is proportional to the shear stress and their surface area (20) (Figure 2A). To quantify the shear force required to remove wildtype cells from the surface, we measured cell departure over a period of 1 minute while simultaneously applying various flow treatments. Specifically, we altered shear rate by changing the flow rate and altered solution viscosity by adding the viscous agent Ficoll to our media. At a shear rate of 4,000 s^-1^ without Ficoll, approximately 0% of wildtype cells departed the surface (Figure 2B). In contrast, at a shear rate of 40,000 s^-1^ without Ficoll, approximately 95% of wildtype cells departed the surface (Figure 2B). As confirmation that shear force drives surface departure, flow at a shear rate of 4,000 s^-1^ with 15% Ficoll (which increases solution viscosity 10x (20)) removed approximately 90% of wildtype cells from the surface (Figure 2B). Together, these experiments demonstrate that a shear force between 10-100 pN is required to remove wildtype cells from a glass surface.

**Figure 2:**
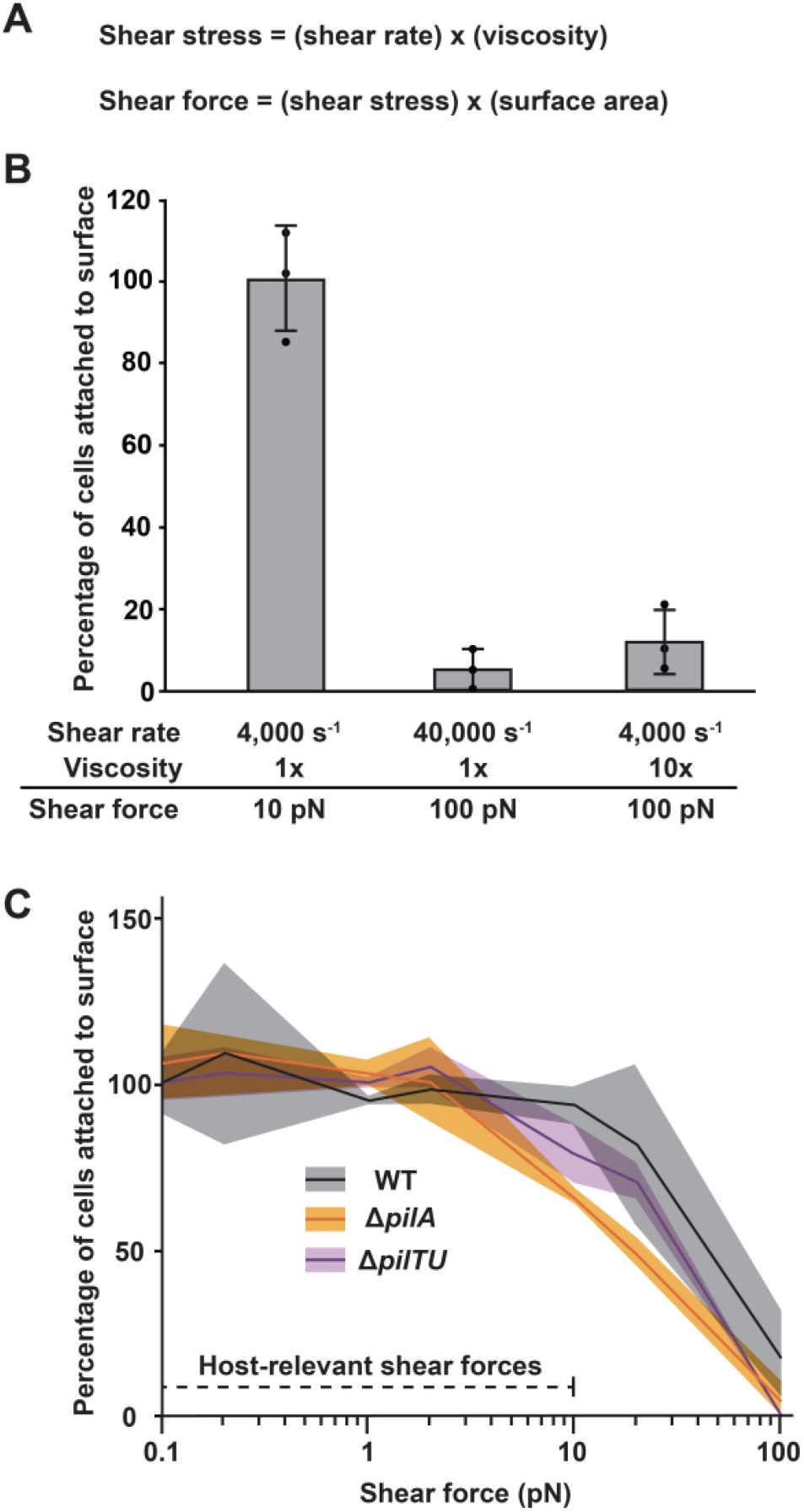
Surface adhered *P. aeruginosa* cells can withstand host-relevant shear forces. (A) In our microfluidic devices, shear stress equals shear rate times fluid viscosity. Shear force is equal to the shear stress times surface area (which we approximate as 2.5 µm^2^). (B) Percentage of WT cells remaining attached to the surface after exposure to 1 minute of different fluid flow treatments. Cells were subjected to different shear forces by varying shear rate and viscosity. Shear rate was modified by changing the flow rate of our syringe pump. 10x viscosity was generated by adding 15% Ficoll, which has been shown previously to modify local viscosity (20, 38). Quantification shows the average and standard deviation of three biological replicates. 4,000 s^-1^ and 40,000 s^-1^ are statistically different with P < 0.01. 4,000 s^-1^ (1x viscosity) and 4,000 s^-1^ (10x viscosity) are also statistically different with P < 0.01. (C) Percentage of cells attached to surface after exposure to 1 minute of fluid flow. WT (black line and gray shading), Δ*pilA* (orange line and shading), and Δ*pilTU* (purple line and shading) cells were subjected to different shear forces, which were generated by varying shear rate. Shading represents the standard deviation of three biological replicates. Total WT cells attached at 0.1 pN is normalized to 100.

To precisely test how flow removes cells from the surface, we subjected wildtype, Δ*pilA*, and Δ*pilTU* cells to a range of shear forces. In host environments such as the bloodstream, urinary tract, and lungs, *P. aeruginosa* cells typically experience shear forces in the range of 0.1 to 10 pN (39–41) (Figure 2C). When subjected to shear forces in the host-relevant regime, wildtype, Δ*pilA*, and Δ*pilTU* cells all remained attached to the surface throughout our experiment (Figure 2C). However, at higher shear forces, flow removed cells. Specifically, a shear force of 100 pN was required to remove the majority of wildtype and Δ*pilTU* cells and a shear force of 20 pN was required to remove the majority of Δ*pilA* cells. As we found no difference between wildtype and Δ*pilTU* cells, the mystery of how Δ*pilTU* cells colonized the surface to greater levels (Figure 1D) remained unanswered. Our observation that very high shear forces were required to remove cells made us question if forcing cells to leave the surface was mechanistically different than allowing cells to depart the surface on their own.

To examine how cells depart the surface on their own, we measured the residence times of cells in a host-relevant flow regime. We defined cell residence time as the time between surface arrival and surface departure (Figure 3A). We quantified the residence times of cells exposed to a shear force of 2 pN as they naturally arrived and departed from the surface (Figure 3A). During the experiment, 77% of wildtype cells had a residence time of less than 2 minutes and 5% had a residence time greater than 10 minutes (Figure 3B). In contrast, Δ*pilA* cells had significantly longer residence times. Specifically, 42% of Δ*pilA* cells resided for less than 2 minutes, while 37% resided for greater than 10 minutes (Figure 3B). The residence times of Δ*pilTU* cells were also significantly longer than wildtype cells, as 69% of Δ*pilTU* resided for less than 2 minutes and 21% of Δ*pilTU* cells resided for greater than 10 minutes (Figure 3B). Together, our results support the conclusion that type IV pili promote surface departure in a host-relevant flow regime. The conclusion that type IV pili promote surface departure provides a rational basis for why Δ*pilTU* cells exhibit an increase in surface colonization (Figure 1D). Furthermore, our observation that type IV pili promote surface departure in a host-relevant flow regime highlights the importance of studying bacteria in mechanically realistic environments.

**Figure 3:**
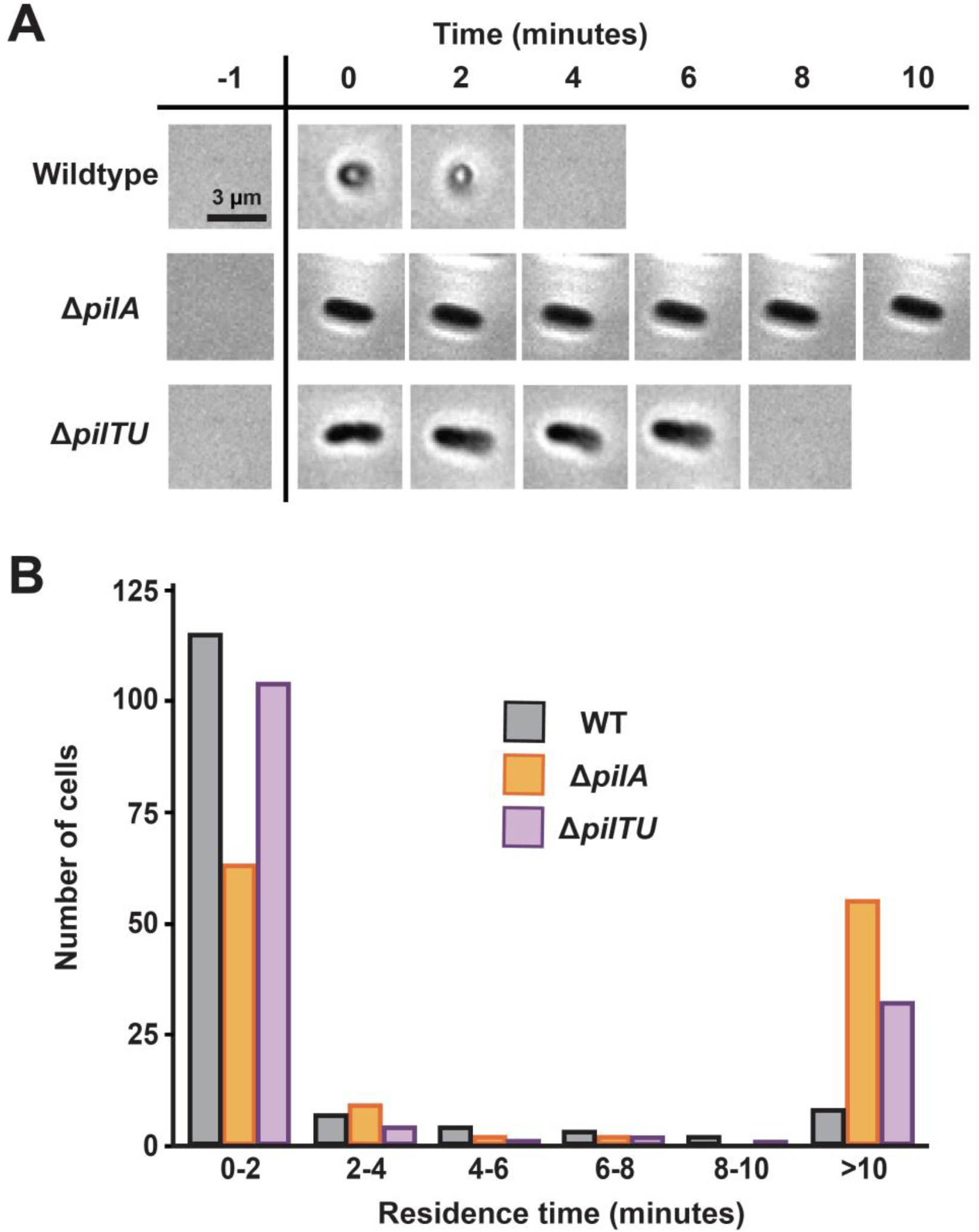
Type IV pili promote *P. aeruginosa* surface departure. (A) Phase images of wildtype, Δ*pilA*, and Δ*pilTU* cells that have arrived on a surface in flow with a shear force of 2 pN over 10 minutes. Images are representative examples for each strain. Scale bar, 3 μm. (B) Surface residence time interval (departure time minus arrival time) of wildtype (WT) (black line and gray shading), Δ*pilA* (orange line and shading), and Δ*pilTU* (purple line and shading) cells in flow with a shear force of 2 pN. 150 cells of each bacterial strain were chosen at random for quantification. Mean residence times of WT and Δ*pilA* are statistically different with P < 0.001. Mean residence times of WT and Δ*pilTU* are statistically different with P = 0.002.

How do type IV pili promote surface departure? A previous report (32, 34) proposed that type IV pili tilt cells off a surface, which subsequently facilitates surface departure. That report focused on *P. aeruginosa* PAO1, a commonly studied isolate of *P. aeruginosa*. As our study is focused on *P. aeruginosa* PA14, the other most commonly studied isolate of *P. aeruginosa*, we explored if surface orientation was altered in our experiments. While the previous study examined bacteria in flow cells, the shear forces used were not in the range of the shear forces in the bloodstream (40), urinary tract (39), and lungs (41). To examine how type IV pili promote surface departure in our experiments, we quantified the cell surface orientation of wildtype, Δ*pilA*, and Δ*pilTU* cells while being simultaneously subjected to host-relevant shear force. To simplify our analyses, we classified cells into three categories: horizontal, tilting, and vertical (Figure 4A). While wildtype cells were approximately evenly distributed between these three categories, Δ*pilA* and Δ*pilTU* cells were mostly horizontal (Figure 4A). Highlighting that type IV pilus retraction promotes cell tilting, 36% of wildtype cells were classified as vertical, while 0% of both Δ*pilA* and Δ*pilTU* cells were vertical (Figure 4A). Thus, our results reinforce previous observations (34) and provide new understanding of how type IV pilus retraction tilts cells away from the surface to promote departure in host-relevant flow environments.

**Figure 4:**
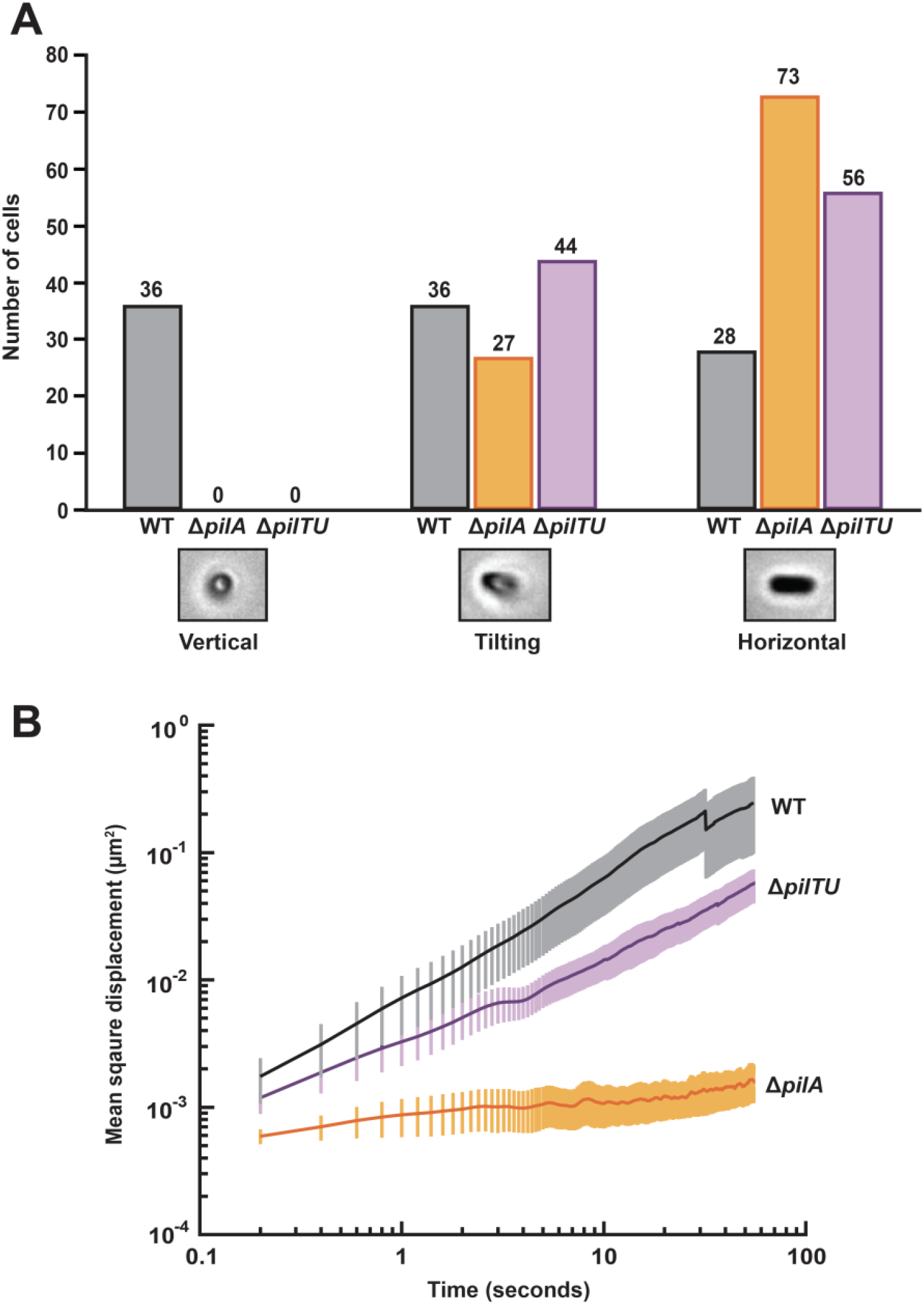
Type IV pili promote *P. aeruginosa* cell tilting and cell migration. (A) Classification of the orientation of cells relative to the surface after exposure to 3 minutes of fluid flow with a shear force of 2 pN. WT (gray bars), Δ*pilA* (orange bars), and Δ*pilTU* (purple bars) cells were manually classified as vertical, tilting, or horizontal based on their appearance in phase images. 100 cells were chosen for classification at random for each bacterial strain. Images show representative examples of each classification. (B) Mean squared displacement (which represents cell motion over time) of WT (black line and gray error bars, n=37 cells), Δ*pilA* (orange line and error bars, n=55 cells), and Δ*pilTU* (purple line and error bars, n=138 cells) cells. Lines represent the average and error bars represent the standard error of the mean. Mean squared displacement was quantified over a 60 second period.

During our experiments, we noticed that Δ*pilA* and Δ*pilTU* mutants exhibited different surface behaviors. For example, the residence times of Δ*pilA* cells were longer than the residence times of Δ*pilTU* cells (Figure 3B). Additionally, the percentage of Δ*pilA* cells that exhibited a horizontal surface orientation was greater than observed with Δ*pilTU* cells (Figure 4A). Based on these observations, we hypothesized that the presence of pili (independent of retraction) changes the adhesive properties of cells on the surface. To test how type IV pili affect the physical interaction between cells and surfaces, we tracked individual cells in flow and measured their mean squared displacement (MSD). MSD represents a physical measurement of how individual cells move relative to their original position over time. Higher MSD values correspond to cells that exhibit more surface movement and migration. As wildtype cells are capable of twitching motility driven by type IV pilus retraction, we hypothesized that wildtype cells would have a high MSD. Our results supported this hypothesis, as wildtype cells had a higher MSD than either Δ*pilA* or Δ*pilTU* mutant strains (Figure 4B). Based on differences we observed between Δ*pilA* and Δ*pilTU* cells, we hypothesized that Δ*pilTU* cells would have a higher MSD than Δ*pilA* cells. In support of that hypothesis, our experiment revealed that MSD values for Δ*pilTU* cells were higher than for Δ*pilA* cells (Figure 4B). Together, these results support the conclusion that the presence of pili (independent of retraction) decrease the adhesive properties of cells on the surface. Furthermore, the pilus-driven decrease in adhesive properties provides an explanation for how wildtype cells depart the surface more frequently than pilus mutants in host-relevant flow contexts.

How does shear force affect the surface residence time of *P. aeruginosa*? A previous study (36) reported the counter-intuitive observation that increasing flow intensity increases surface residence time of *P. aeruginosa* PA14. To confirm this observation, we quantified surface residence time of wildtype cells exposed to shear rates of 160 s^-1^ or 1,600 s^-1^ (Figure S4). In support of the previous study, we observed that only 6% of wildtype cells resided for greater than 10 minutes at 160 s^-1^ (Figure 5B & S4). In contrast, when subjected to a shear rate of 1,600 s^-1^, 37% of wildtype cells resided for greater than 10 minutes (Figure S4). The previous study suggested that the shear force associated with flow was responsible for the increase in surface residence time. As shear force is proportional to shear rate and solution viscosity, we explicitly tested the hypothesis that shear force increases residence time by using the viscous agent Ficoll. When cells were exposed to a shear rate of 160 s^-1^ without Ficoll, 76% of wildtype cells resided for less than 2 minutes (Figure 5B & S4). However, at a shear rate of 160 s^-1^ with 15% Ficoll (which generated 10 times more shear force (20)), only 15% of wildtype cells resided for less than 2 minutes (Figure 5B). Collectively, our results explicitly demonstrate that increasing shear force generates the counter-intuitive outcome of enhancing surface adhesion.

**Figure 5:**
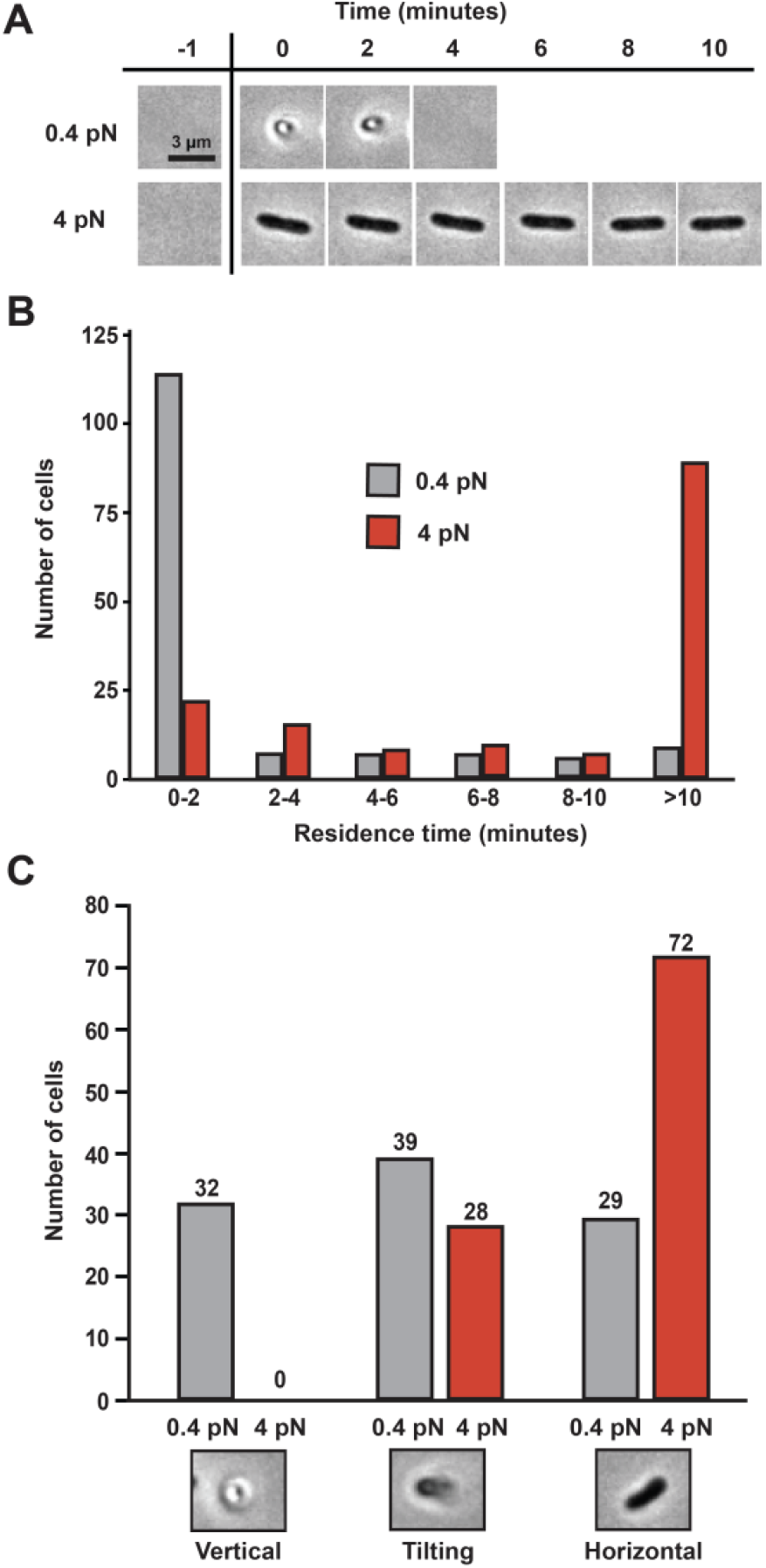
Shear force enhances *P. aeruginosa* adhesion by counteracting cell tilting. (A) Phase images of wildtype cells that have arrived on a surface in flow with a shear force of 0.4 pN or 4 pN over 10 minutes. A shear force of 0.4 pN was generated by a flow with a shear rate of 160 sec^-1^ with 0% Ficoll, and a shear force of 4 pN was generated by a flow with a shear rate of 160 sec^-1^ with 15% Ficoll. Images are representative examples for each strain. Scale bar, 3 μm. (B) Surface residence time interval (departure time minus arrival time) of WT cells in flow with a shear force of 0.4 pN (gray bars) or 4 pN (red bars). 150 cells were chosen at random for each flow condition for quantification. Mean residence times of WT cells subjected to 0.4 pN and 4 pN are statistically different with P < 0.001. (C) Classification of cell-surface orientation of wildtype cells after exposure to 3 minutes of fluid flow with a shear force of 0.4 pN (gray bars) or 4 pN (red bars). A shear force of 0.4 pN was generated by a flow with a shear rate of 160 sec^-1^ with 0% Ficoll, and a shear force of 4 pN was generated by a flow with a shear rate of 160 sec^-1^ with 15% Ficoll. Cells were manually classified as vertical, tilting, or horizontal based on their appearance in phase images. 100 cells were chosen for classification at random for each flow condition. Images show representative examples of each classification.

While the mechanism by which shear force enhances adhesion of *P. aeruginosa* is unknown, there are multiple theoretical explanations for this phenomenon. For example, shear-enhanced adhesion of *E. coli* is mediated by the adhesin FimH, which uses a catch bond that resembles a finger trap. It is also possible that shear force could enhance adhesion by changing the geometric orientation of rod-shaped cells to increase surface contact. Based on our results, we favored this explanation and hypothesized that shear force enhances adhesion by promoting horizontal cell attachment. To test how flow intensity affects surface orientation, we quantified the cell surface orientation of wildtype cells exposed to shear rates of 160 s^-1^ or 1,600 s^-1^ (Figure S5). We observed that wildtype cells exposed to a shear rate of 160 s^-1^ were approximately evenly distributed between our three orientation categories (Figure 5C, S5). However, the majority of wildtype cells exposed to a shear rate of 1,600 s^-1^ were found in the tilting or horizontal orientation (Figure S5). To explicitly test how shear force affects surface orientation, we altered solution viscosity by using the viscous agent Ficoll. When exposed to a shear rate of 160 s^-1^ with 15% Ficoll, we found that wildtype cells were predominantly in the tilting and horizontal orientations (Figure 5C). These results demonstrate that shear force tips cell over, promotes horizontal cell orientation, and decreases the frequency of surface departure.

To precisely examine how shear force affects cell surface association, we tracked individual cells exposed to different flow conditions. First, we aimed to test if increasing shear rate would lower the MSD of cells, hypothesizing that more flow would result in firmer surface association. In support of our first hypothesis, cells exposed to a shear rate of 1,600 s^-1^ had a lower MSD than cells exposed to shear rate of 160 s^-1^ (Figure S5, S6). Second, we aimed to test if increasing solution viscosity would lower the MSD of cells, hypothesizing that more shear force would result in firmer surface association. In support of our second hypothesis, cells exposed to a shear force of 4 pN had a lower MSD than cells exposed to shear force of 0.4 pN (Figure S6, S7). Together, these experiments provide evidence that shear force restricts the motion of cells on a surface, which likely promotes *P. aeruginosa* colonization in host-relevant flow regimes.

## Discussion

Our experiments reveal how shear force and type IV pili coordinate *P. aeruginosa* surface departure. Using a Δ*pilA* mutant, we showed that type IV pili have an important role in cell surface arrival and surface colonization in flow (Figure 1). Additionally, we demonstrated that type IV pili have an important role in cell surface departure in host-relevant flow regimes (Figure 3). Furthermore, we discovered that Δ*pilTU* cells (which lack pilus retraction) are better than wildtype at colonizing a surface (Figure 1), which can be explained by the increased residence time of Δ*pilTU* cells (Figure 3). By independently modifying shear rate and viscosity, we determined that while very high shear forces promote surface departure (Figure 2), host-relevant shear forces prevent surface departure (Figure 5). Through precise cell tracking, we established that the cell surface orientation angle and cell MSD are associated with increased surface departure (Figure 4, Figure 6). Together, our results highlight how physical and biological factors combine to drive surface departure of the human pathogen *P. aeruginosa*.

For many years, type IV pili have been recognized as important factors driving bacterial surface arrival (2, 23, 24, 37). However, more recent reports have suggested that type IV pili also have an important role in bacterial surface departure (32–34). Our data supports that idea that type IV pili drive both surface arrival and departure. As type IV pili extend away from the cell body and make direct contact with the surface, it is intuitive to understand how they promote surface arrival. However, the mechanism by which type IV pili drive surface departure may be less obvious. We conclude that type IV pili promote surface departure in two ways. First, we conclude that pilus retraction leads to the movement and tilting of cells off the surface, which increases the likelihood of their complete dissociation from the surface. This conclusion is supported by our data that the residence time of Δ*pilTU* cells is longer than wildtype (Figure 3). Second, we conclude that the presence of pili on the cell body impacts the physical association of cells with the surface in such a way that cells are less firmly attached, experience more movement, and are more likely to dissociate from the surface. This conclusion is supported by the fact that the residence time of Δ*pilA* cells is longer than Δ*pilTU* cells (Figure 3). Together, our data highlight how type IV pili orchestrate the dynamics of *P. aeruginosa* surface interactions in flow.

Intuitively, one would expect that shear force removes adherent cells from a surface. However, we show that shear force can enhance surface adhesion. Conceptually, there are multiple mechanisms that could allow for force-enhanced adhesion (6–14). For example, a surface protein could utilize a conformational change similar to a children’s toy known as a finger trap which shrinks the circumference of a tube when pulling force is applied. In fact, the FimH adhesin from *E. coli* uses a finger trap-like mechanism to strengthen its adhesive properties when subjected to shear force (10). Another mechanism that could allow for force-enhanced adhesion involves reorientation and establishment of new adhesive contacts (10). The data we have presented suggests that this mechanism underlies the force-enhanced adhesion of *P. aeruginosa*. Specifically, our data show that shear force reorients cells with respect to the surface (Figure 5), creating a situation where new adhesive contacts can form. Our MSD data shows decreased surface motion when cells are reoriented by shear force (Figure 4, Figure 5), supporting the interpretation that shear force enhances the association between cells and the surface. Based on findings presented here, it is clear that the human pathogen *P. aeruginosa* optimizes surface association in response to host-relevant shear force, which potentially explains its impressive ability to colonize a diverse array of host niches.

How much shear force do cells experience during infection? For an organism like *P. aeruginosa* which infects many sites of the human body, there are a wide range of shear forces it may experience. For example, the bloodstream typically generates shear forces ranging from 0.1 pN to 10 pN (40), the lung typically generates shear forces ranging of 1 pN to 5 pN (41), and the urinary tract typically generates shear forces ranging from 0.1 pN to 5 pN (39). As *P. aeruginosa* cells are likely to experience 0.1 to 10 pN of shear force during host colonization, it is logical to hypothesize that they have evolved mechanisms to optimize surface association when exposed to host-relevant shear forces. Our data strongly support this hypothesis, as wildtype cells adhere well when subjected to shear forces between 0.1 and 10 pN and are only removed from the surface when subjected to 100 pN (Figure 2). Based off multiple pieces of evidence, our central conclusion is that *P. aeruginosa* optimizes surface adhesion to promote surface colonization in flow-rich host environments. In support of our central conclusion, orthogonal approaches (42) have recently generated a comprehensive mathematical model that explains how *P. aeruginosa* uses type IV pili to optimize the adhesion-migration trade-off during colonization. Thus, there is a growing body of evidence (42) that *P. aeruginosa* couples type IV pilus dynamics and cell geometry to optimize surface adhesion and cell migration in fluctuating mechanical environments.

## Acknowledgements

We thank Anuradha Sharma, Alex Shuppara, Gilberto Padron, Nick Martin, Lisa Wiltbank, Vada Becker, Ben Bratton, Dan Kearns, Ankur Dalia, Paola Mera, and Thomas Kehl-Fie for helpful discussions and comments on the manuscript. This work was supported by funding from the European Research Council through a starting grant for B.S. (BacForce, g.a. No. 852585). This work was also supported by start-up funds from the University of Illinois at Urbana-Champaign and grant K22AI151263 from the National Institutes of Health to J.E.S.

## Contributions

J.S.P., A.N.S, B.S., and J.E.S. designed research. J.S.P. performed research. J.S.P., A.N.S., and J.E.S. analyzed data. J.S.P. and J.E.S. wrote the paper.

## Supplementary Information for

## Materials and Methods

### Bacterial strains, plasmids, and growth conditions

Bacterial strains used in this study are described in Table 1 the plasmids used are described in Table 2, and the primers used are described in Table 3. *P. aeruginosa* strains were grown on LB agar plates (1.5% Bacto Agar) and in liquid LB in a roller drum at 37°C. LB medium was prepared using premix Miller LB Broth (BD Biosciences) and using standard LB preparation protocols.

**Table 1:** *P. aeruginosa* strains used in this study.

| *P. aeruginosa* strain | Description | Source |
| --- | --- | --- |
| PA14 | wildtype; clinical isolate from burn wound | (43) |
| JS177 | $\Delta pilA::aacC1$ | This paper |
| JS176 | $\Delta pilA::FRT$ | This paper |
| JS183 | $\Delta pilTU::aacC1$ | This paper |
| JS182 | $\Delta pilTU::FRT$ | This paper |
| JS215 | $Tn7::PpilA-pilA::aacC1$ | This paper |
| JS218 | $Tn7::PpilA-pilA::FRT$ | This paper |
| JS217 | $Tn7::PpilT-pilTU::aacC1$ | This paper |
| JS220 | $Tn7::PpilT-pilTU::FRT$ | This paper |

**Table 2:** Plasmids used in this study.

| Plasmid | Description | Source |
| --- | --- | --- |
| pAS03D | Plasmid to generate deletion mutants in *P. aeruginosa* PA14 | (44) |
| pFLP2 | Plasmid expressing FLP2 to recombine FRT sites | (45) |
| pUCP18-RedS | Lambda red recombineering vector | (44) |

**Table 3:**
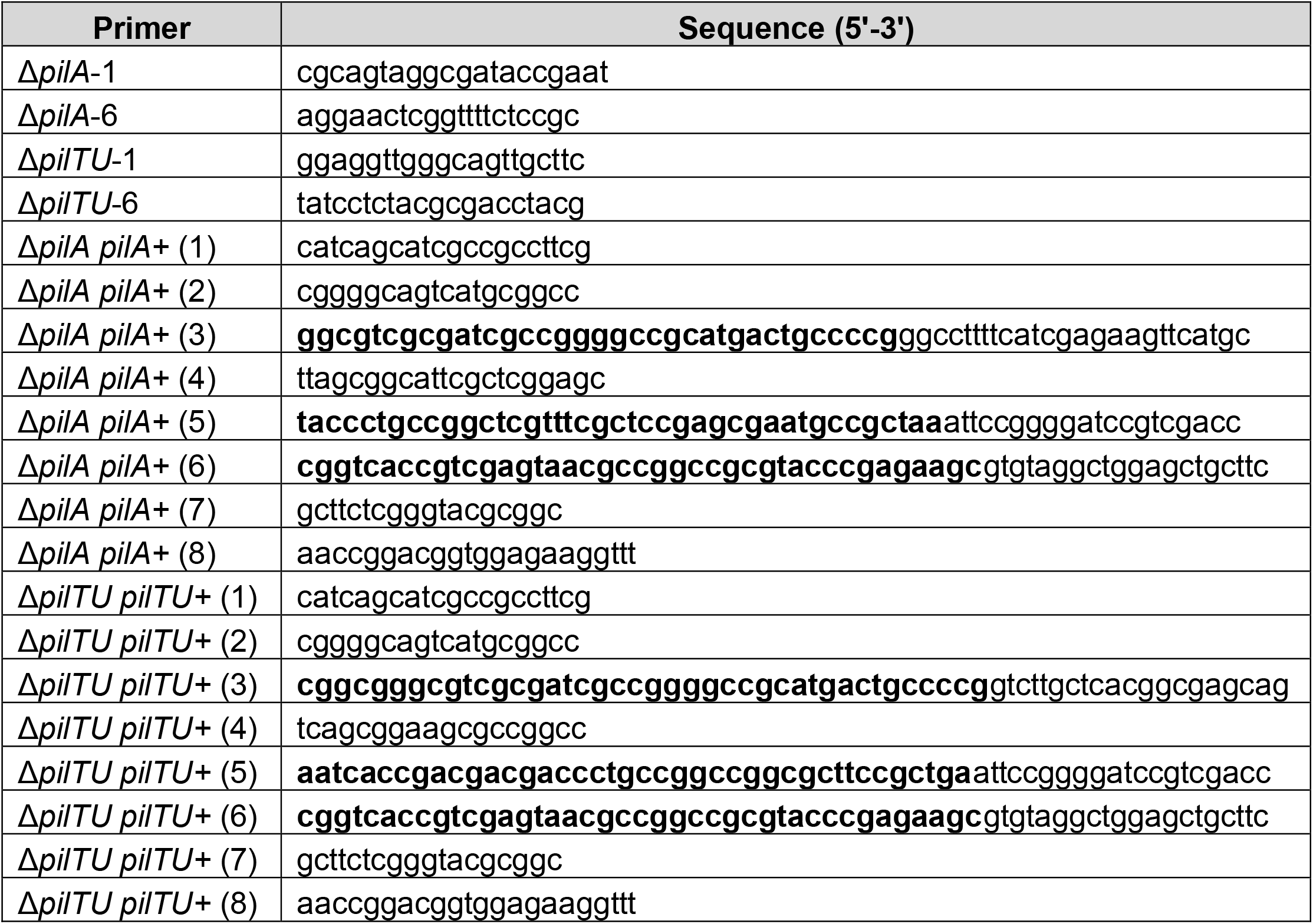
Primers used in this study.

#### Generation of *P. aeruginosa* mutants

Gene deletions were generated using the lambda Red recombinase system. The deletion construct was Gibson-assembled from three PCR products. First fragment, approximately 500 bp upstream of the target insertion site was amplified from PA14 genomic DNA. Second fragment, containing *aacC1* ORF flanked by FRT sites was amplified from pAS03D. Third fragment, approximately 500 bp downstream of the target insertion site was amplified from PA14 genomic DNA. The Gibson-assembled product was transformed into PA14 cells expressing the plasmid pUCP18-RedS. The colonies were selected on 30 μg/ml gentamycin, the mutants of interest were counter-selected on 5% sucrose, and pFLP2 was used to flip out the antibiotic resistance gene. pUCP18-RedS and pFLP2 were selected for using 300 μg/ml carbenicillin.

#### Construction of microfluidic devices

Microfluidic devices were created using soft lithography techniques. Devices were designed on Illustrator (Adobe Creative Suite) and masks were printed by CAD/Art Services. Molds were made on 100 mm silicon wafers (University Wafer) and spin coated with SU-8 3050 photoresist (MicroChem). Polydimethylsiloxane (PDMS) chips were plasma-treated to bond with glass slides for at least 24 hours prior to experiments. The devices used in all experiments contained 7 parallel channels (500 μm width x 50 μm height x 2 cm length). Channels individually contained an inlet tube and an outlet tube. PDMS chips were plasma treated to a 60 mm x 35 mm x 0.16 mm superslip micro cover glass (Ted Pella, Inc.).

#### Colonization and residence time assays

*P. aeruginosa* cells were loaded into plastic 5 mL syringes (BD) at mid-log phase with an optical density of approximately 0.5. All microfluidic experiments were performed at ∼ 22 °C. The device set-up involves the loaded syringes attached to tubing which connects the needle to the inlet of the device (BD Intramedic Polyethylene Tubing; 0.38 mm inside diameter, 1.09 mm outside diameter). These syringes were situated on a syringe pump (KD Scientific Legato 210) which was used to produce fluid flow. The outlet of the device employed the same tubing and vacated into a bleach-containing waste container. The syringe pump was used to generate flow rates of 10 μL/min, which correspond to shear rates of 800 s^-1^.

#### Surface departure assay

Microfluidic channels were seeded with *P. aeruginosa* cells at mid-log phase with an optical density of approximately 0.5. All microfluidic experiments were performed at ∼ 22 °C. Cells were injected into the microfluidic device using a pipette and were allowed to attach on the glass surface for 10 minutes prior to exposure to flow. The device set-up involves the use of plastic 5 mL syringes (BD) with attached tubing connecting the needle to the inlet of the device (BD Intramedic Polyethylene Tubing; 0.38 mm inside diameter, 1.09 mm outside diameter). These syringes were situated on a syringe pump (KD Scientific Legato 210) which was used to produce fluid flow. The outlet of the device employed the same tubing and vacated into a bleach-containing waste container. The syringe pump was used to generate flow rates of 0.5-500 μL/min, which correspond to shear rates of 40 s^-1^ – 40,000 s^-1^.

#### Phase contrast microscopy

Images were obtained with a Nikon Ti2-E microscope controlled by NIS Elements. All images were taken with Nikon 100x Plan Apo Ph3 1.45 NA objective, a Hamamatsu Orca-105 Flash4.0LT+ camera, and Lumencor Sola Light Engine LED light source.

#### Tracking of cell motion and mean squared displacement

Bacterial cells were tracked from phase-contrast movies using the ImageJ plugin MicrobeJ. MicrobeJ automatically identifies the cells in each frame, segments the frames, and connects the recognized cells from all frames. The cell segmentation and tracking are then corrected and connected manually via the ImageJ GUI. Bacterial cells that are not in complete view during the tracking were excluded from the analysis. The cells were tracked starting from the time of arrival on the surface, until the cell departed from the surface. If one pole of the cell is detached and the other pole of the cell remained attached, the tracking of that cell still occurs.

**Figure S1:**
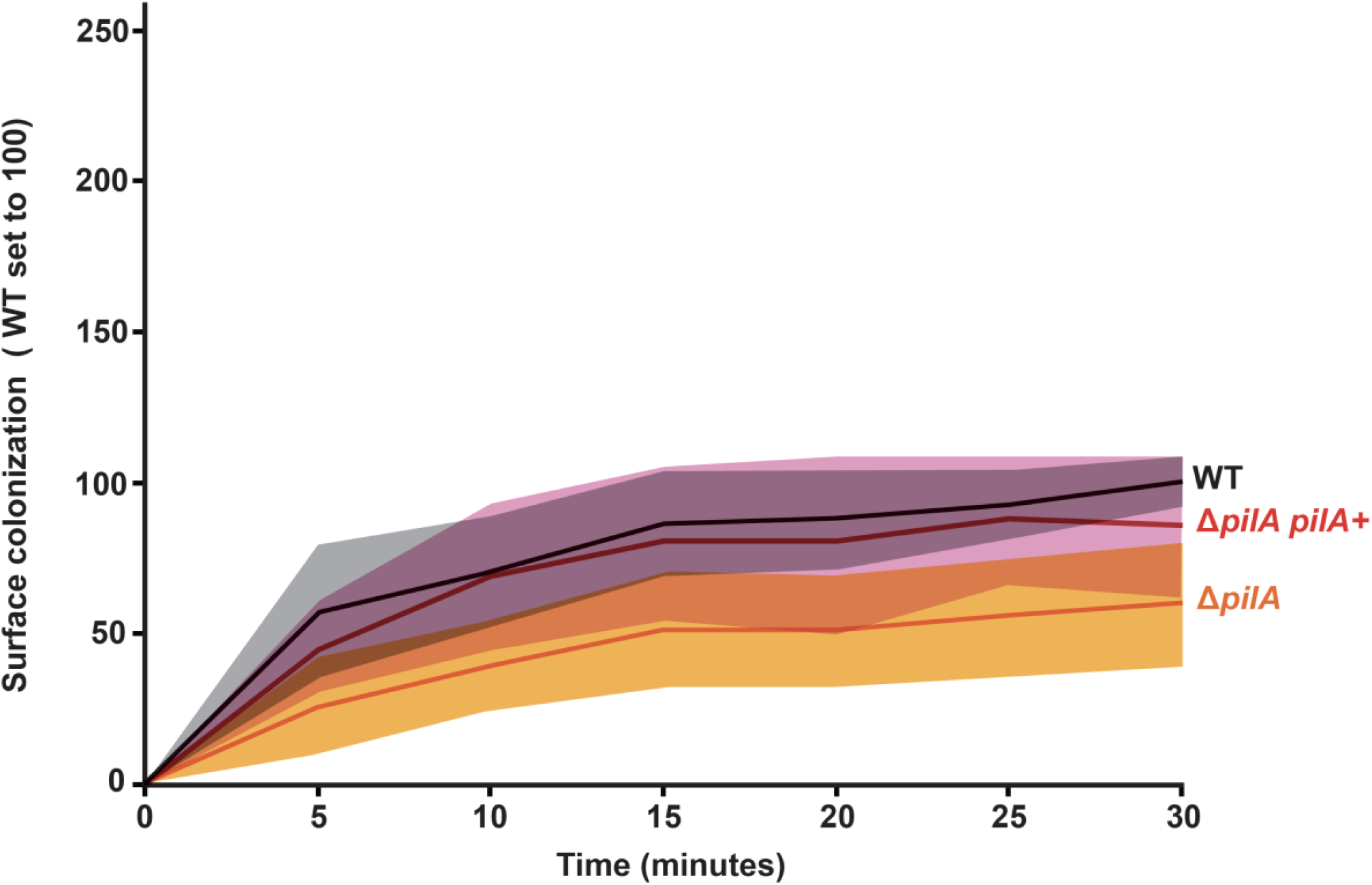
Re-introduction of *pilA* complements the Δ*pilA* surface colonization phenotype. Surface colonization of an empty channel with cells flowing in over 30 minutes. WT (black line and gray shading), Δ*pilA* (orange line and shading), and Δ*pilA pilA+* (red line and pink shading) cells were introduced at a shear rate of 800 sec^-1^. WT cell surface accumulation at 30 minutes is normalized to 100. Shading represents the standard deviation of three biological replicates.

**Figure S2:**
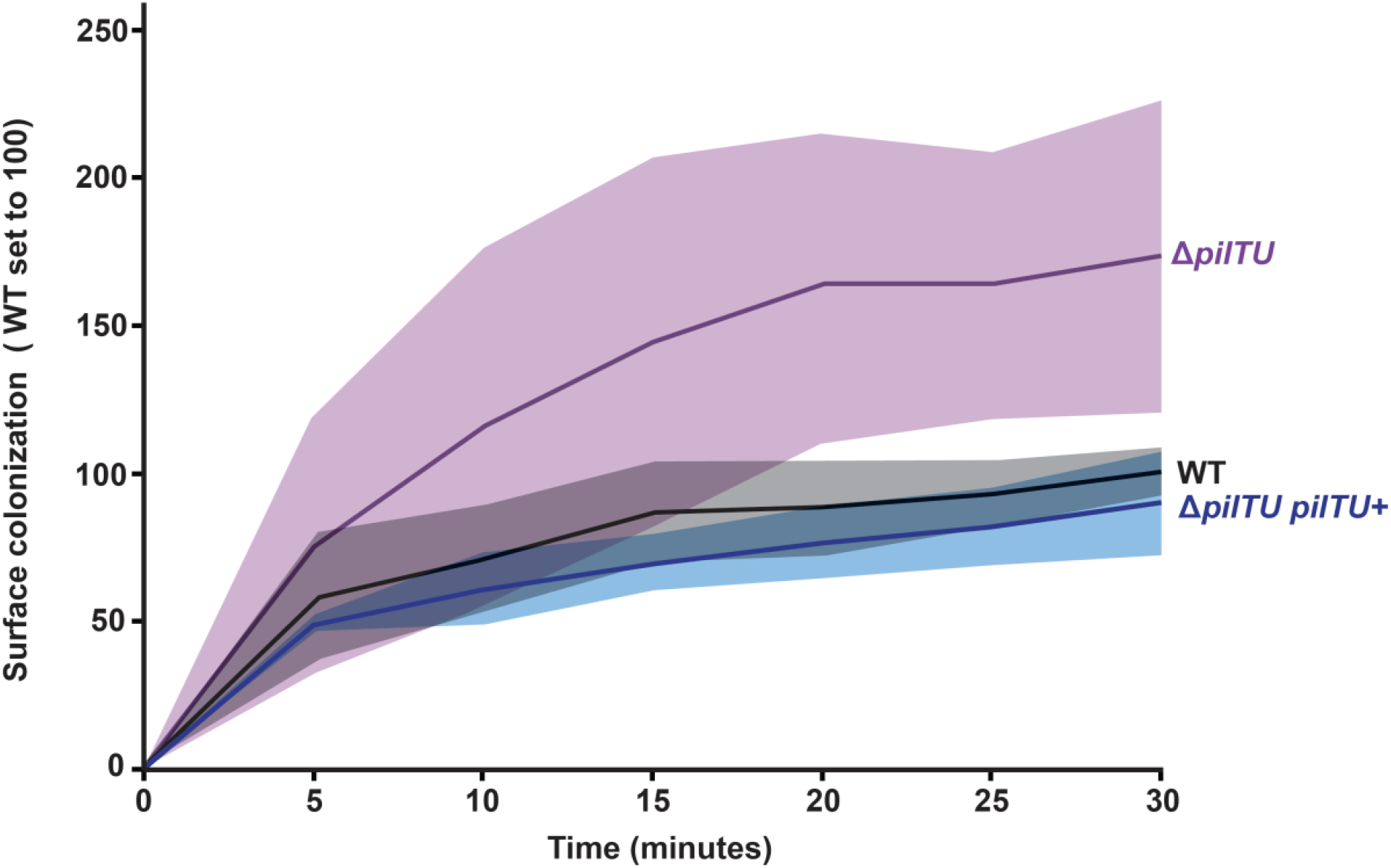
Re-introduction of *pilTU* complements the Δ*pilTU* surface colonization phenotype. Surface colonization of an empty channel with cells flowing in over 30 minutes. WT (black line and gray shading), Δ*pilTU* (purple line and shading), and Δ*pilTU pilTU+* (blue line and shading) cells were introduced at a shear rate of 800 sec^-1^. WT cell surface accumulation at 30 minutes is normalized to 100. Shading represents the standard deviation of three biological replicates.

**Figure S3:**
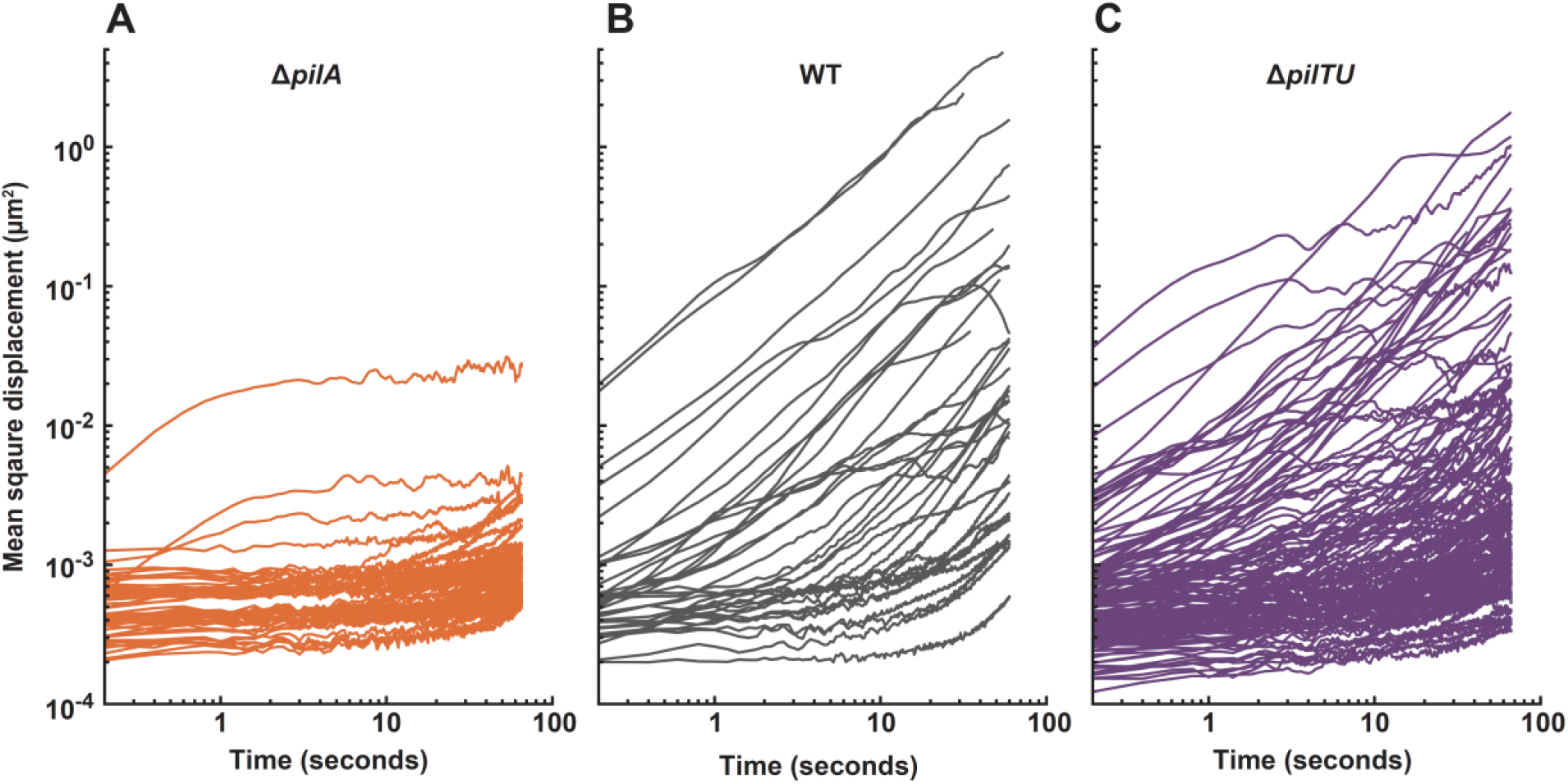
Type IV pili promote *P. aeruginosa* cell migration. Individual cell trajectory traces of Δ*pilA* (orange lines), WT (gray lines), and Δ*pilTU* (purple lines). WT (gray lines, n=37), Δ*pilA* (orange lines, n=55), and Δ*pilTU* (purple lines, n=138).

**Figure S4:**
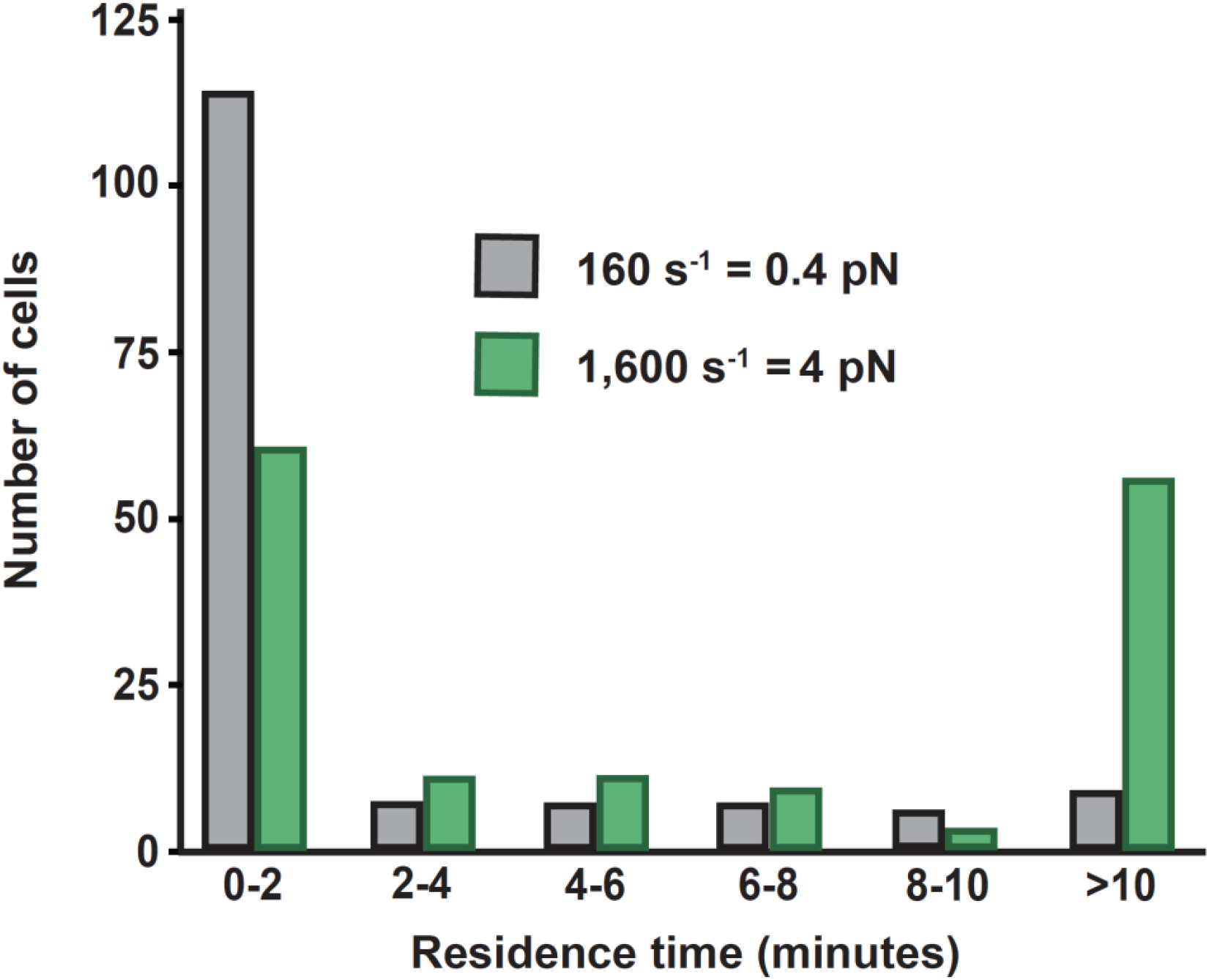
Shear force enhances *P. aeruginosa* adhesion. Residence time of wildtype cells attached to surface after exposure to varying shear forces for 30 minutes. Wildtype cells subjected to shear force of 160 sec^-1^ (gray bars) and wildtype cells exposed to shear force of 1,600 sec^-1^ (green bars). 150 cells were chosen at random for each flow condition for quantification. Mean residence times of WT cells subjected to 160 sec^-1^ and 1,600 sec^-1^ are statistically different with P < 0.001.

**Figure S5:**
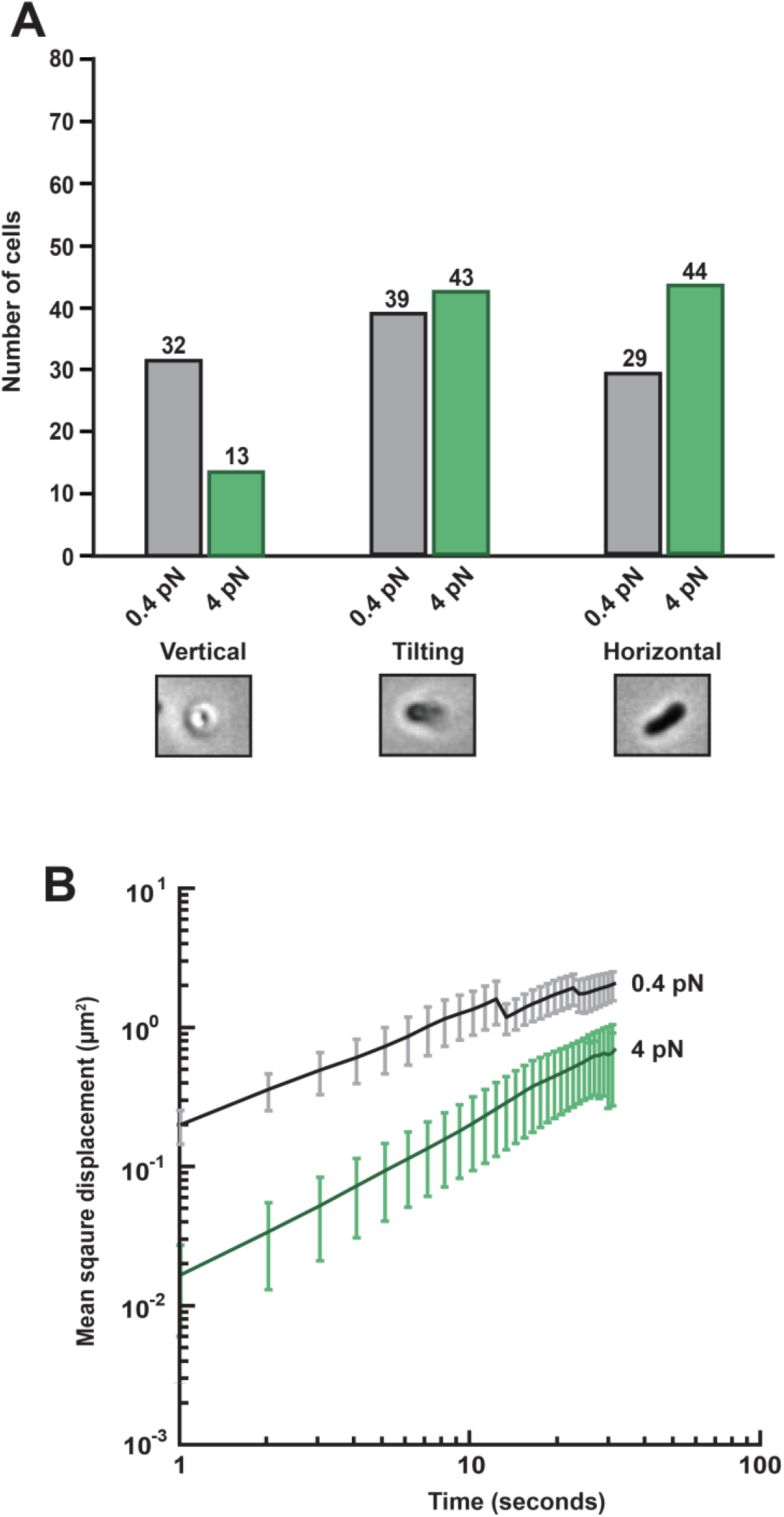
Shear force counteracts *P. aeruginosa* cell tilting and cell migration. (A) Quantification of cell attachment orientation to surface after exposure to 3 minutes of fluid flow. Wildtype cells subjected to shear force of 160 sec^-1^ (gray bars) and wildtype cells exposed to shear force of 1600 sec^-1^ (green bars). Images represent surface orientation of cells. (B) Average mean squared displacement (cell trajectory) of wildtype cells under two conditions. Wildtype cells subjected to shear force of 160 sec^-1^ (black line with gray shading, n=24) and wildtype cells exposed to shear force of 1600 sec^-1^ (green line and shading, n=24).

**Figure S6:**
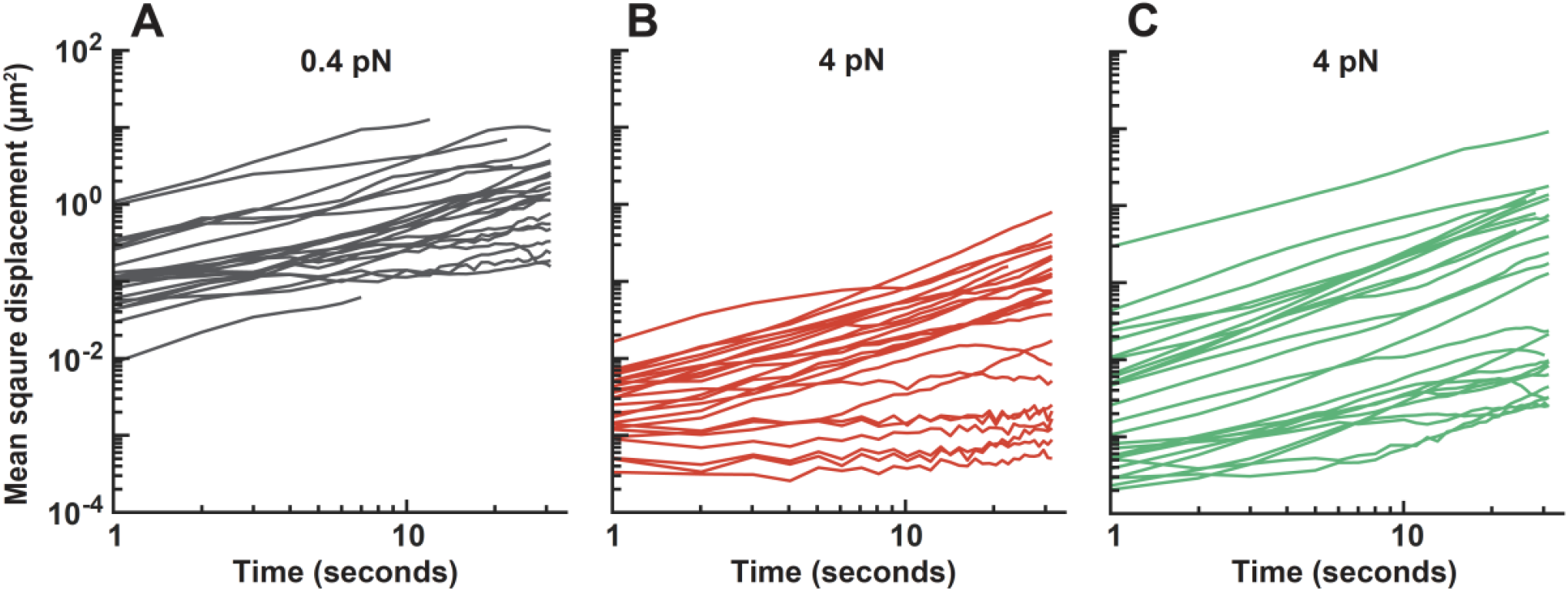
Shear force restricts *P. aeruginosa* cell migration in a host-relevant flow regime. Individual cell trajectory traces of WT cells under three conditions. WT cells subjected to shear forces of 160 sec^-1^ (A, gray lines, n=24), 160 sec^-1^ with 15% Ficoll (B, red lines, n=24), and 1600 sec^-1^ (C, green lines, n=24).

**Figure S7:**
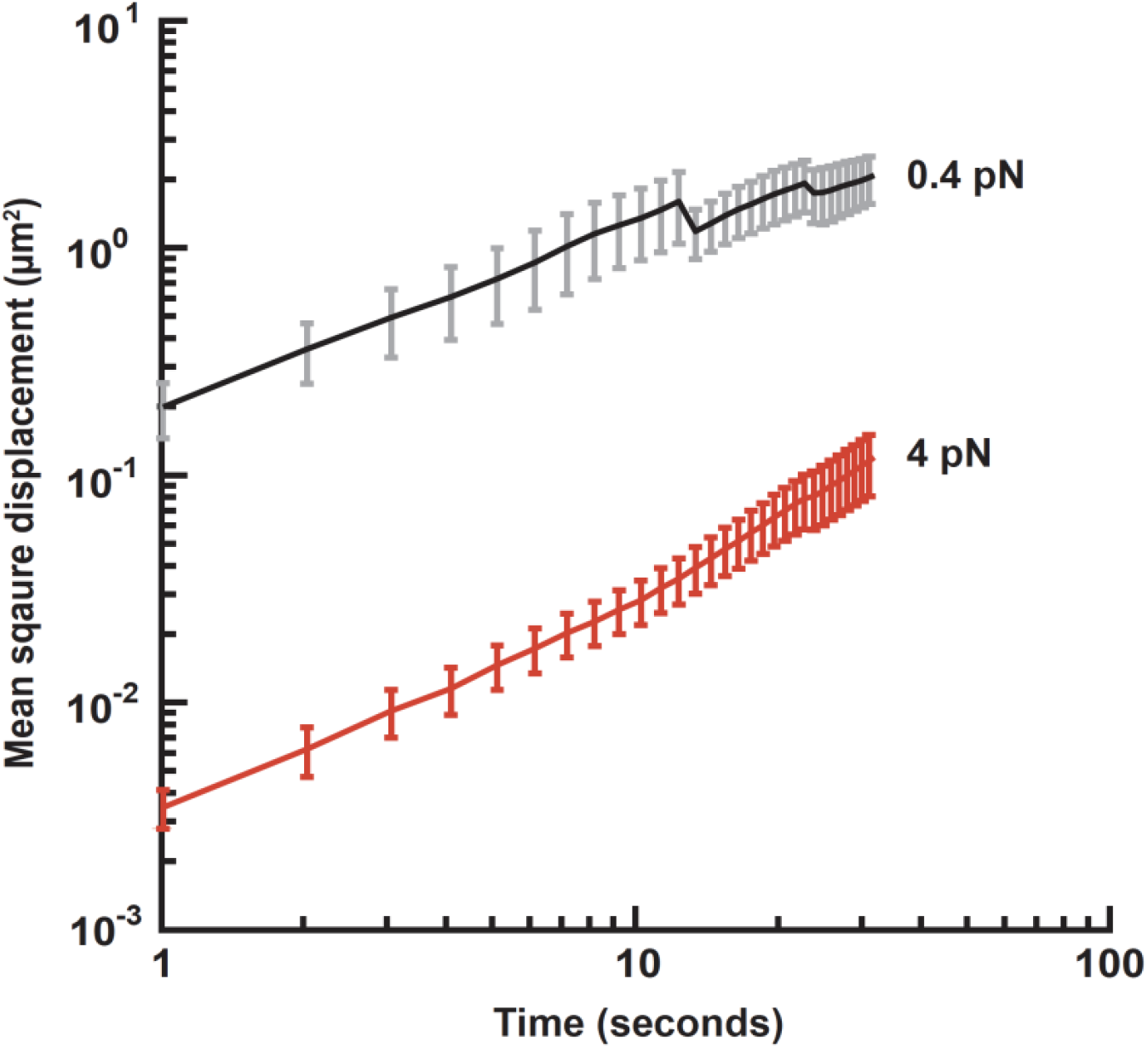
Shear force restricts *P. aeruginosa* cell migration in a host-relevant flow regime. Mean squared displacement (which represents cell motion over time) of wildtype cells attached to the surface experiencing a shear force of 0.4 pN (black line with gray error bars, n=24 cells) or 4 pN (red line and error bars, n=24 cells). Lines represent the average and error bars represent the standard error of the mean. Mean squared displacement was quantified over a 30 second period.

